# Structural and functional analysis of the human Cone-rod homeobox transcription factor

**DOI:** 10.1101/2022.01.09.475544

**Authors:** Penelope-Marie B. Clanor, Christine Buchholz, Jonathan E. Hayes, Michael A. Friedman, Andrew M. White, Ray A. Enke, Christopher E. Berndsen

## Abstract

The cone-rod homeobox (CRX) protein is a critical K50 homeodomain transcription factor responsible for the differentiation and maintenance of photoreceptor neurons in the vertebrate retina. Mutant alleles in the human gene encoding CRX result in a variety of distinct blinding retinopathies, including retinitis pigmentosa, cone-rod dystrophy, and Leber congenital amaurosis. Despite the success of using in vitro biochemistry, animal models, and genomics approaches to study this clinically relevant transcription factor over the past 24 years since its initial characterization, there are no high-resolution structures in the published literature for the CRX protein. In this study, we use bioinformatic approaches and small-angle x-ray scattering (SAXS) structural analysis to further understand the biochemical complexity of the human CRX homeodomain (CRX-HD). We find that the CRX-HD is a compact, globular monomer in solution that can specifically bind functional cis-regulatory elements encoded upstream of retina -specific genes. This study presents the first structural analysis of CRX, paving the way for a new approach to studying the biochemistry of this protein and its disease-causing mutant protein variants.

## Introduction

The vertebrate retina is a multilayered neural sensory tissue located in the posterior portion of the eye. Visual perception occurs when photons of light enter the front of the eye, are focused onto the retina, and are then absorbed by rod and cone photoreceptors (PR), initiating the biochemical process of phototransduction ^1^. Several key transcription factors regulate the development, differentiation, and maintenance of these critical PR neurons during retinogenesis and in the mature retina ^2^.

TFs are proteins that modulate the transcriptional activity of genes. TFs directly affect the differentiation and maintenance of diverse cell types in various tissues by controlling transcriptional networks. Homeodomain TFs contain a homeodomain (HD), a globular domain of about 60 amino acids conferring the biochemical ability to bind specific DNA motifs ^3^. Paired-type HD TFs recognize a TAAT core DNA motif which is further specified by the amino acid at position 50 of the HD ^3^. This sequence specificity can be binned into distinct subclasses. HDs with a glutamine residue at position 50 (Q50 type) preferentially bind a TAAT<u>TA/G-</u>motif, while those with a lysine residue at position 50 (K50 type) favor a TAAT<u>CC</u> motif ^4–6^. HD family proteins represent 15-30% of TFs in plants and animals and regulate diverse developmental processes ^7^.

Cone rod homeobox (CRX) is a critical K50 HD TF for regulating PR and bipolar cell (BC) transcriptional networks during differentiation and cell fate determination in the vertebrate retina ^2,8^. CRX functions by binding to TAATC cis-regulatory elements (CREs) encoded in genomic DNA sequences in and adjacent to PR-specific genes regulating their transcription ^9,10^. CRX binding to CREs is mediated by the highly conserved N-terminal HD DNA binding domain ^11^. Transcriptional regulation is achieved by an array of protein-protein interactions mediated by CRX’s activation domain (AD) encompassing the C-terminal half of the protein (Figure 1A) ^11,12^.

**Figure 1:**
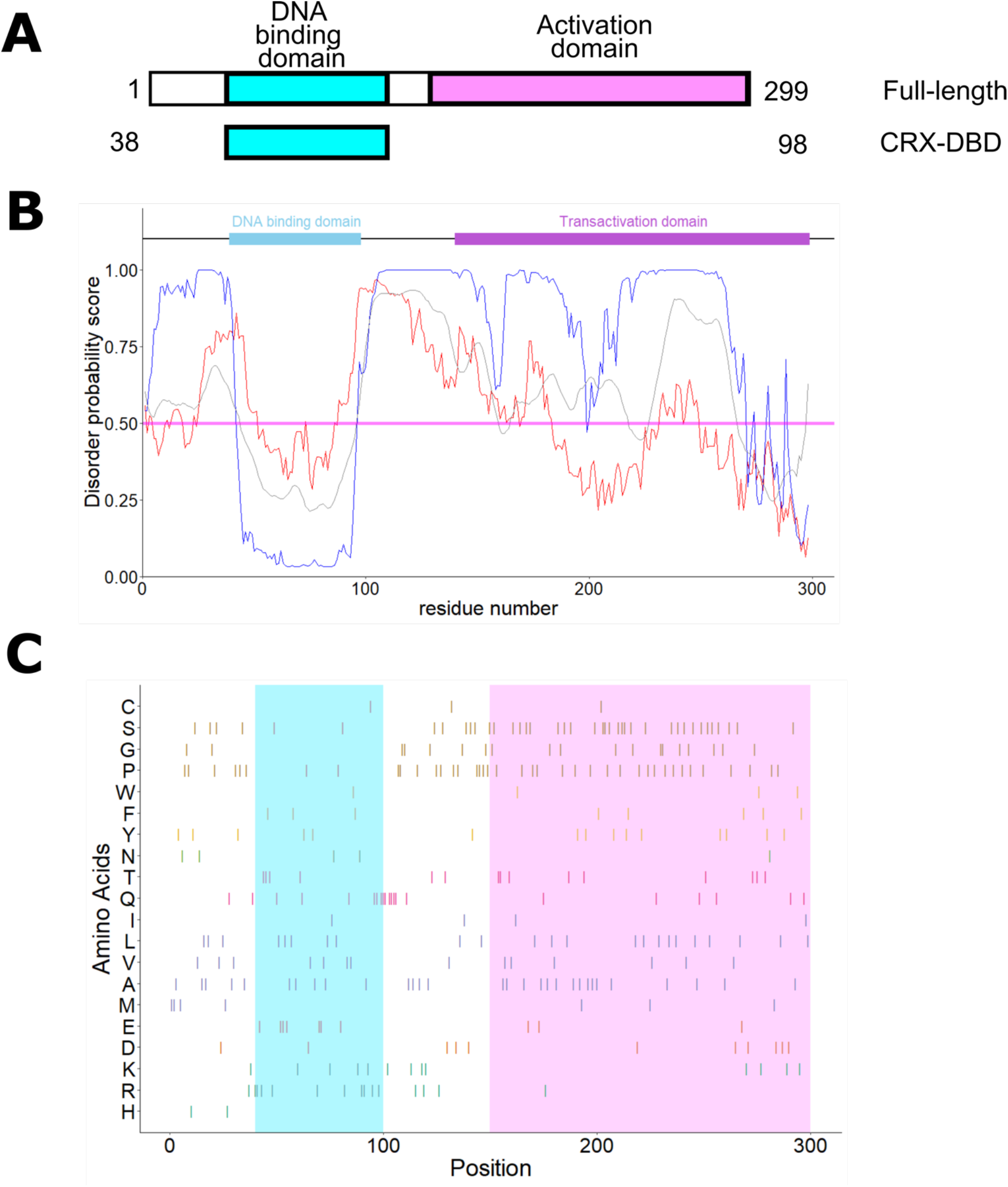
Human CRX protein organization and sequence analysis. (**A**) Cartoon showing the domain structure of human CRX (referred to as full-length CRX) and the DNA binding domain of CRX (CRX-DBD). Numbers indicate the amino acids positions encompassed in each construct. (**B**) Disorder prediction of the full-length CRX sequence. The data from IUPred2A shown in red, from ODIN shown in blue, and from PrDOS in grey (see methods for details). (**C**) Amino acid prevalence within the CRX sequence. The cyan box indicates the DNA binding domain and the salmon box indicates the activation domain.

Numerous clinically relevant mutations in the *CRX* gene result in an array of biochemical dysfunction and they are responsible for multiple blinding retinopathies, including retinitis pigmentosa (RP), cone-rod dystrophy (CoRD), and Leber congenital amaurosis (LCA) ^13^. Due to the complexity of the CRX protein, these mutant alleles can be categorized into subclasses based on allele type, biochemical dysfunction of the protein, and resulting pathology ^14^. While animal models and in vitro experiments have helped characterize the various mechanisms of CRX-induced pathology, there are currently no high-resolution NMR or X-ray structures in the published literature for the CRX protein to aid in this endeavor. In this study, we use small-angle x-ray scattering (SAXS) to analyze the three-dimensional properties of the human CRX DNA binding HD protein. This study presents the first structural analysis of this clinically relevant protein and opens the door to a new modality for studying CRX and its disease-causing mutant alleles.

## Methods

### Reagents and peptides

All buffers and chemicals were ordered from Fisher Life Science or Sigma-Aldrich. A peptide corresponding to the human CRX homeodomain (amino acids 39 to 98) was synthesized by Genscript.

### Computational Sequence Analysis

DNA and protein sequences of human CRX were obtained from Uniprot entry O43186.

For the sequence properties analysis in Figures 1B and C, the protein sequence was imported into R, and the amino acids were binned by properties. For visualization of the pathogenic substitutions, data were downloaded from a search of CRX from ClinVar ^15^. In R, substitutions linked solely to CRX and which were described as pathogenic or likely pathogenic and either missense, premature stop, frameshift substitutions were identified. Data were then plotted using the ggplot2 package.

The primary sequence of CRX was provided to PrDOS ^16^, IUPred2A ^17^, and ODiNPred ^18^ servers for disorder prediction. The resulting predictions were plotted using the ggplot2 package. CRX binding regions (CBRs) obtained from a previously published human retina ChIP-seq study ^10^ were visualized in the UCSC Genome Browser, hg38 human genome assembly, and were used to identify putative CRX binding motifs in the proximal promoter regions of the human *RHO* and *PDE6B* genes.

### CRX Cloning & Expression

The coding sequence for the N-terminal 107 amino acids comprising the DNA binding domain (DBD) of the human CRX protein was fused to a 6x His-tag inserted following the met start codon and subcloned into the commercial PMAL-c5x expression plasmid containing a maltose binding domain (MBD) coding sequence using Xmnl and EcoR1 restriction sites (GenScript, Piscataway, NJ). CRX DBD-MBD plasmid was transformed into BL21 E. coli competent cells and grown on LB plates supplemented with ampicillin for selection overnight at 37°C. Transformant colonies were used to inoculate 10 mL starter cultures followed by 1L LB cultures grown to an optical density of 0.5-0.7 OD at 600 nm at which time they were induced with 0.2 mM IPTG and cultured at 37 °C for 1 hour at 200 RPM. Cultures were pelleted, resuspended in lysis buffer, and disrupted using three rounds of probe sonication at 75% amplitude for five second on/off pulses with one minute between pulse sequences. Lysates were pelleted and the soluble fraction was used for downstream protein purification steps.

### Protein Purification

The 6X-His-CRX-DBD-MBD fusion protein was purified from lysates using a multistep chromatography strategy on an AKTA Start FPLC. Initial purification of the MBD portion of the fusion protein was facilitated using amylose chromatography. The 1mL MBPTrap™ HP column was equilibrated with 20 mM Tris-HCl, 200 mM NaCl, 1 mM EDTA (pH 7.4) binding buffer and protein was eluted with binding buffer with an additional 10 mM maltose. Fractions containing CRX-DBD-MBD, identified by SDS-PAGE and FPLC UV spectra, were pooled.

Heparin column chromatography was used to facilitate the removal of contaminating bacterial nucleic acids from the fusion protein. The 1 mL HiTrap™ column was equilibrated with 0.1M sodium phosphate (pH 7.4) and protein eluted with 10 mM sodium phosphate, 2 M NaCl using a linear gradient yielding 72 column volume (pH ∼7). Size exclusion chromatography was used as a final step to separate heavier and lighter molecular weight contaminating proteins from the desired protein. The Hiload™ 16/600 Superdex™ 200 pg column was equilibrated with 0.02 M NaPO_4_, 50mM NaCl (pH 7). All FPLC fractions were collected and examined with SDS PAGE and 260nm/280nm ratio to determine purity, quality, and quantity of the final purification product.

### Small Angle X-ray Scattering (SAXS) analysis

The CRX DBD peptide was resuspended in water before being dialyzed overnight at 4 °C into 50 mM sodium phosphate, pH 7, 100 mM NaCl, and 5 mM imidazole. Samples were shipped overnight at 4 °C to the SIBYLS beamline at the Advanced Light Source. Samples were injected into an Agilent 1260 series HPLC with a Shodex KW-802.5 analytical column at a flow rate of 0.5 ml/min. Small-angle X-ray scattering (SAXS) data was collected on the elution as it came off of the column. The incident light wavelength was 1.03 Å at a sample to detector distance of 1.5 m. This setup results in scattering vectors, q, ranging from 0.013 Å^-1^ to 0.5 Å^-1^, where the scattering vector is defined as q = 4πsinθ/λ and 2θ is the measured scattering angle.

Radially averaged data were processed in SCATTER (ver 4.0d) and RAW (ver. 2.1.1) to remove the scattering of the sample solvent and identify peak scattering frames ^19^. RAW was used to calculate the dimensions of the molecule, the molecular weight, and the pair-distance distribution function (PDDF). Plots of data were produced in R using the ggplot2 package ^20^.

### *ab initio* modeling of CRX shape

The electron density of the peptide was calculated in RAW using the DENSS program^21^.

DENSS predicted 51 shapes followed by averaging and refinement. The SAXS data were also used in the DAMMIF program in RAW to create a dummy atom model ^22^. The DAMMIF program created 15 models followed by refinement and averaging. The results from DENSS and DAMMIF were each visualized in the program CHIMERA in order to produce a mesh model from each ^23^. Each of these models were then individually fitted with a homology model created from the DBD sequence in the program trRosetta ^24^.

### Circular dichroism

For CD measurements, the cuvette contained CRX-DBD at a concentration of 24 µM in 50 mM sodium phosphate, 100 mM sodium chloride, and 5 mM imidazole in a 0.2 cm quartz cuvette. Data were collected from 320 nm to 200 nm at 1 nm/sec with 3 accumulation on a Jasco J-1500 CD spectrometer. Spectra were collected at temperatures from 10-95 °C at an interval of 5 °C, with a gradient of 2°/min. The sample was equilibrated at each temperature for 1 min before spectra were collected. Data were exported in .csv formatted and plotted in R using the ggplot2 package. Melting temperatures were calculated through a global fit to a one-step irreversible unfolding model using the Calfitter app ^25^.

### DNA binding assays

35 bp double-stranded DNA oligonucleotide probes corresponding to putative CRX binding sites within the human *RHO* and *PDE6B* proximal promoter CBRs were commercially synthesized (Figure 4C). Corresponding point mutant probes were designed for each by swapping a thymine base, critical for CRX affinity, for an adenine base (gaTta to gaAta). 35 ng of dsDNA probes were combined with 300 ng of the human CRX-DBD-MBD fusion protein in 1X PBS for 1 hour at 37 °C and visualized on a 1% agarose gel post stained with GelRed.

## Results

### The CRX activation domain is predicted to be flexible and disordered

The K50 homeodomain transcription factor CRX has been previously shown to orchestrate complex transcriptional regulation in photoreceptor retinal neurons; however, the detailed biochemical mechanism and structural detail of how this critical protein stimulates these effects are not yet apparent. CRX binds to CREs via a conserved N-terminal HD and imparts transcriptional regulation mediated by C-terminal AD (Figure 1A). Previously, we modeled the CRX DNA binding domain (DBD) structure, suggesting that it formed a compact helical structure similar to other homeodomain-containing transcription factors ^26^. Building on this analysis, we assessed the viability of modeling the structure of the transactivation domains of CRX. Unlike the CRX DBD, this portion of CRX shows little homology to known protein structures, and nearly 75% of the protein is predicted to be disordered (Figure 1B). However, we found some disagreement between the prediction methods, especially in the C-terminal AD (Figure 1B). The recent AlphaFold prediction of the proteomes for model species contains entries for human, mouse, rat, and zebrafish CRX, and all of the entries show higher predicted error outside of the DBD ^27,28^. Within amino acid positions 150 to 299 of the activation domain are 50 prolines and serines, which is 34% of the amino acid composition of the total protein (Figure 1C). The high number of proline residues, which is known to disrupt the secondary structure, and serine, which only moderately favors helical structures, likely leads to the apparent flexibility and disorder in this region ^29^. The DBD is rich in basic amino acids, which allows the protein to interact with the backbone and bases of the DNA. Thus, we could not confidently model the structure of any region of CRX outside of the DBD and expand our previous model.

### Solution shape of the CRX DNA binding domain

Given the limited structural information on CRX, we attempted to crystallize the homeodomain of CRX, but we have been thus far unsuccessful. Concomitantly, we pursued studies of the structure through small-angle X-ray scattering (SAXS), which is useful for determining the size, shape, and molecular weight of biomolecules ^30^. To obtain a homogeneous sample, we fractionated a synthesized peptide of the human CRX homeodomain (CRX-DBD) via size exclusion chromatography. The protein eluted as a single peak and we calculated the molecular weight to be 6.4 kDa with the expected molecular weight from the sequence being 7.4 kDa (Figure 2A). The molecular weight is consistent with this protein being monomeric in solution. We next performed circular dichroism on the peptide to confirm that the peptide was folded. From measurement at 222 nm, we calculated that the CRX-DBD melted at 57.4 °C (95% C.I. 2.5 °C;) (Figure 2B). We next submitted the peptide for size exclusion chromatography coupled to SAXS detection. This approach was chosen in part because we observed that the CRX-DBD peptide had stability and solubility issues in equilibrium SAXS experiments (data not shown). From our data collection, we observed that the CRX-DBD peptide migrated as a single peak and we were able to average six frames to get a data set (Figure 3A). We calculated an R_g_ of 11.8 ± 0.2 with a molecular weight of 5.7 kDa (98.7% C.I. 5.0 to 6.0 kDa), in line with our molecular mass value calculated from an independent size exclusion chromatography experiment. The dimensionless Kratky plot for CRX-DBD peptide shows a single peak with a descent toward the x-axis characteristic of a globular protein structure in solution (Figure 3B).

**Figure 2:**
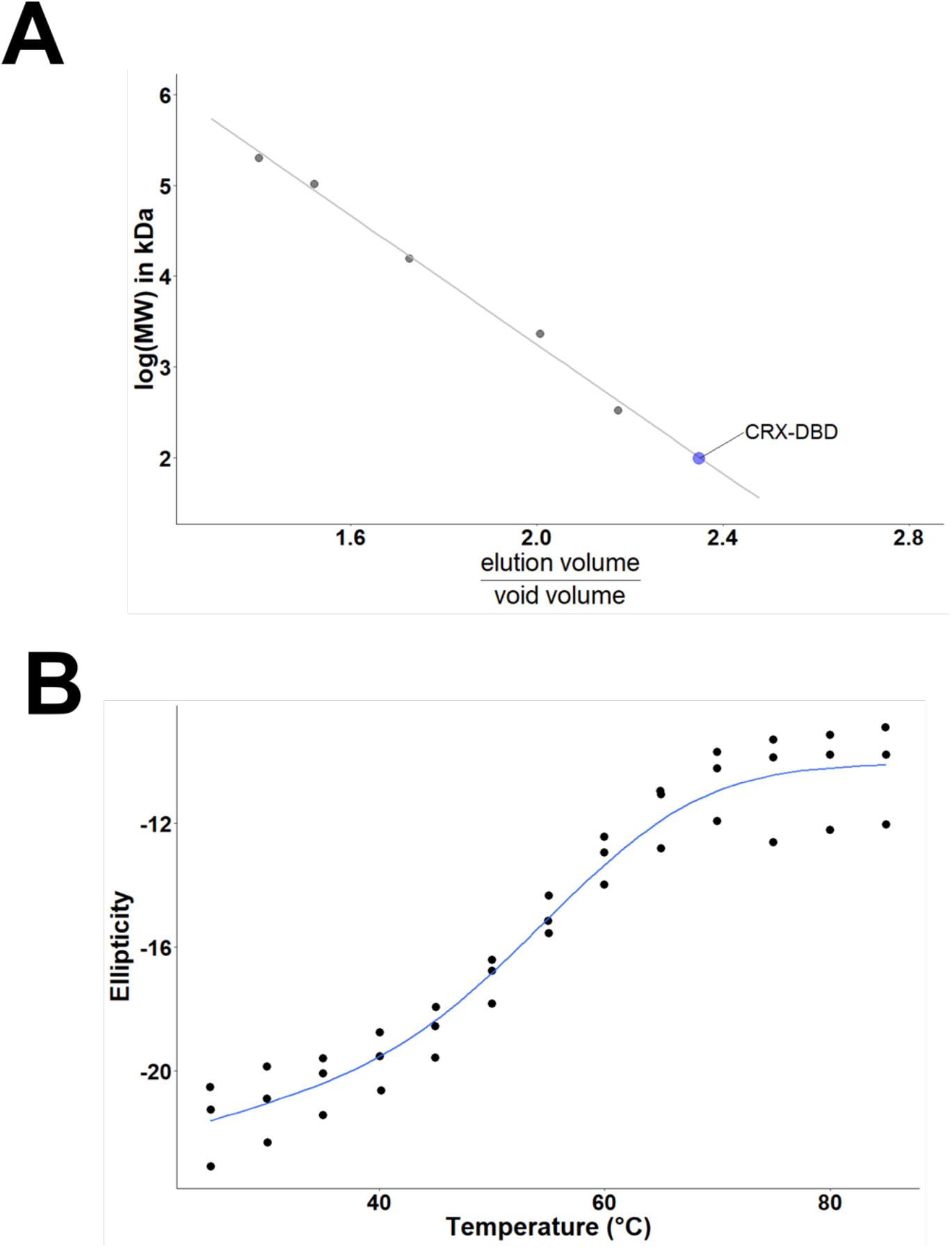
Solution properties of the CRX-DBD peptide. (**A**) Elution of CRX-DBD from the size exclusion column. The peptide eluted as a single peak with a calculated molecular weight of 6.3 kDa. These data are consistent with a CRX-DBD monomer. (**B**) Melting curve of CRX-DBD at 222 nm. The calculated melting temperature was 57.4 Three separate experiments are shown on the plot along with the fitting line from CalFitter.

**Figure 3:**
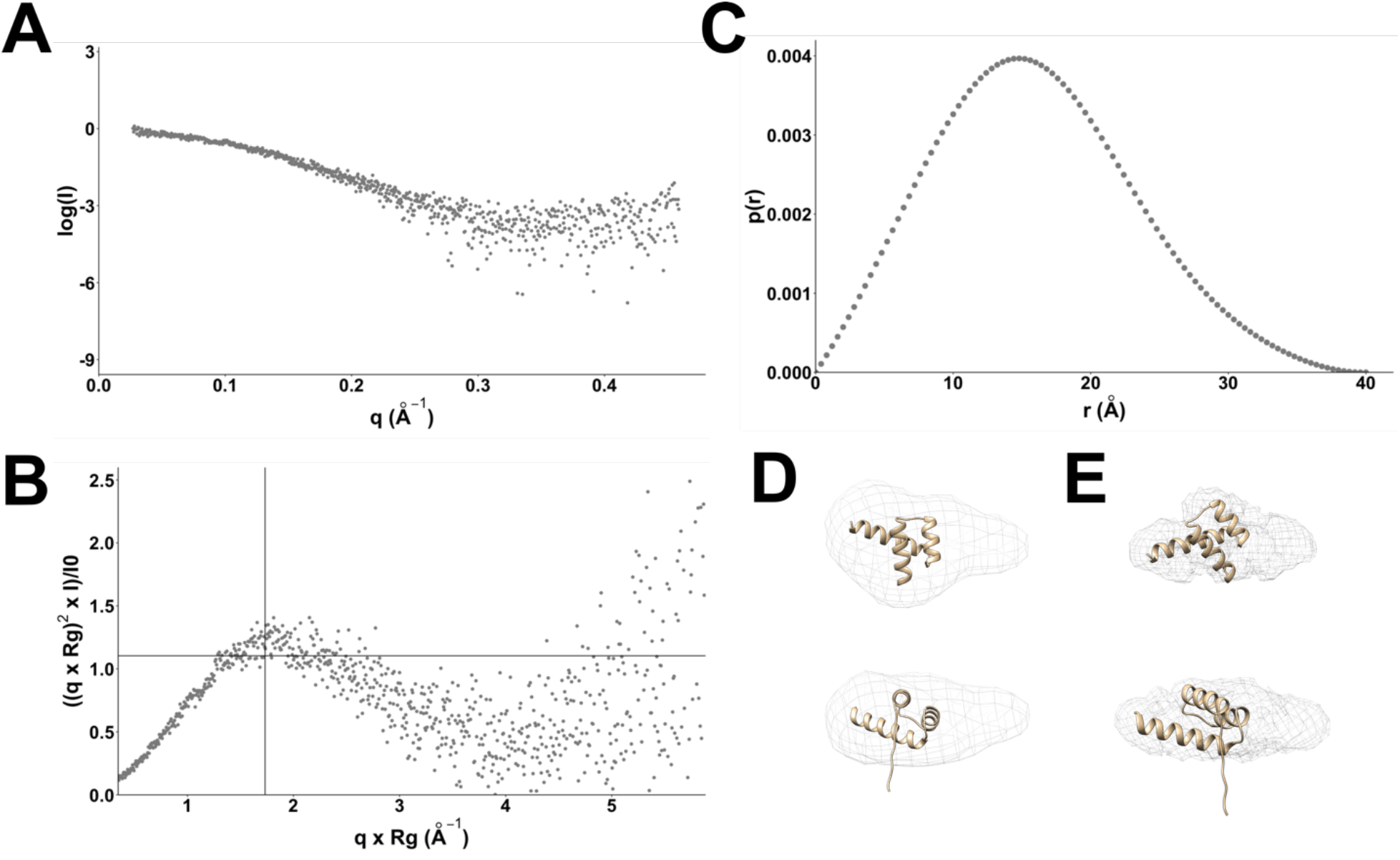
Structural analysis of CRX-DBD. (**A**) log of intensity vs momentum transfer for the averaged frames containing CRX-DBD (**B**) Kratky analysis of CRX-DBD (**C**) Pair-distance distribution plot (**D**) envelope of the dummy atom model produced by DAMMIF aligned to the CRX-DBD homology model (**E**) Electron density map produced from DENSS aligned to the CRX-DBD homology model.

We then fitted the SAXS data using DAMMIF and DENSS to predict the 3-D shape.

DAMMIF simulates the overall shape from the SAXS data by calculating several bead models and then selecting the ones that fit best to further refine and fit the data ^22^. DENSS uses iterative structure factor retrieval to calculate the 3-D shape and electron density ^21^. From these analyses, the envelope resolution calculated in DENSS was 17 and the ensemble resolution calculated in DAMMIF was 22 ± 2 Å. The DAMMIF program calculated the mean normalized spatial discrepancy (NSD) of the model to be 0.61 ± 0.04. We fitted these models to the trRosetta homology model to determine the accuracy of the model (Figures 3D and 3E). The homology model fits the envelope from the DAMMIF fitting and the electron density from DENSS. The

DBD model contains an N-terminal region that does not have a defined structure. It is possible that this region could adopt several conformations that could occupy the empty portions of the density (Figures 3D and 3E). These data further support our findings that the CRX-DBD forms a monomer in solution and has a globular structure.

### The CRX-DBD binds wild-type but not mutated CREs in the human genome

We next analyze the ability of the CRX-DBD to bind to cis-regulatory elements (CREs) in the human genome using a MBP tagged CRX-DBD fusion protein in mobility shift assays. The MBP tag enhances the shift in the migration of DNA upon binding to the DBD, which aids in resolving bands. Recent ChIP-seq analysis of CRX binding in the genome of human retinal neurons demonstrates CRX binding regions (CBRs) upstream of the photoreceptor-specific genes *RHO* and *PDE6B* (Figures 4A and 4B)^10^. Further sequence analysis of these regions revealed 5’-GATTA-3’ (5’-TAATC-3’ on reverse strand) putative CRX binding motifs present within each CBR (Figure 4C). The *RHO* upstream region contains two motifs separated by a 4 bp spacer whereas the *PDE6B* upstream region contains a single motif. To determine if these specific motifs within CBRs are functional CREs, we synthesized 35 bp dsDNA oligonucleotides corresponding to the wt sequences as well as point mutant oligos introducing a single base change at a key motif residue (Figure 4C). Incubation of the CRX-DBD-MBD fusion protein with wt *RHO* and *PDE6B* oligos induced a gel shift indicating both regions contained functional CRX CREs (Figures 4D and 4E). To determine if either of the *RHO* motifs are functional CREs, substitutions of the critical 3rd position of each motif were made. The gel shift was abolished with a base substitution in the first motif, but persisted with a substitution in the second motif, indicating CRE functionality of only one of these motifs (Figure 4D). A similar analysis of the single PDE6B motif demonstrates its functionality as a bonafide CRE (Figure4E). Collectively, these data demonstrate the functionality of the CRX-DBD protein used in this study. These findings also suggest that CRX binding affinity is dictated by features other than the local base-pair grammar of binding motifs.

**Figure 4:**
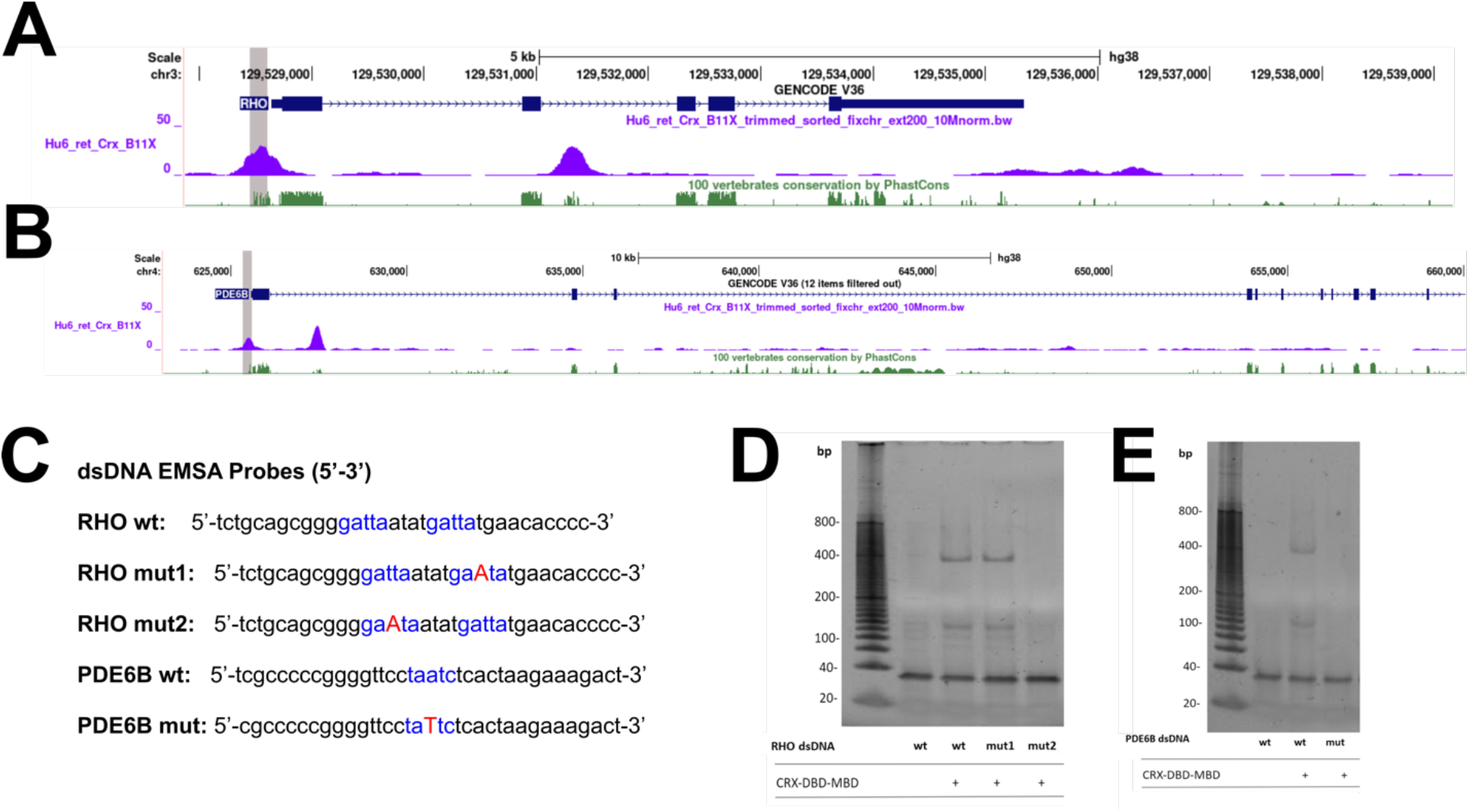
CRX binds evolutionarily conserved CREs in the human genome. UCSC genome browser outputs of the human (**A**) *RHO* and (**B**) *PDE6B* loci aligned with BAM density plots from an adult human retina CRX ChIP-seq experiment identify CRX binding regions (CRBs) in both loci. Sequences highly conserved between vertebrate species are indicated as green peaks in the PhastCons Conservation data track. (**C**) Sequences of dsDNA probes used for EMSAs in the *RHO* and *PDE6B* promoter regions. Five base pair predicted CRX motifs within CBRs are indicated in blue lettering. Mutated bases are shown in red and capital letters to indicate the position and identity of the change. EMSA assays for (**D**) the *RHO* promoter and (**E**) the *PDE6B* promoter. 34ng of was used for each assay, and 300 ng purified human MBP-CRX-DBD protein added to lanes indicated with +.

## Conclusions/Discussion

The retina-specific K50 homeodomain transcription factor CRX is required for the differentiation and maturation of photoreceptor PR neurons in the mammalian retina. While the biological functions of CRX are well described, a detailed exploration of its structural biochemistry is less clear. To address this knowledge gap, we characterized the structure of the CRX DNA binding domain and its binding to functional DNA CREs. Our data show that the CRX-DBD is monomeric in solution, adopts a compact structure, and is able to bind to specific DNA motifs.

Homeodomain proteins exist in a variety of quaternary structures, including homodimers.

In contrast, our size exclusion chromatography and SAXS data clearly show that the CRX homeodomain is monomeric in solution (Figure 2A). We do observe two bands in the DNA binding assays, which could be consistent with a DNA-CRX-DBD dimer and a DNA-CRX-DBD ternary complex. Conversely, Chen and coworkers showed previously that DNA sequences with two CRX binding sites likely bound one CRX per site while those with a single site did not show behavior consistent with a CRX dimer ^31^. This study did use a longer section of CRX in their studies, which may indicate that dimerization occurs outside of the 60 amino acids that we consider to be the DNA binding domain in the present work ^11,31^. However, the precise species observed in the present assays is not clear, and further structural and biochemical work is required.

We further found that a substitution in the third position of the motif abolished binding (Figures 4D and 4E). However, when two motifs were present, only the first motif was necessary for CRX binding (Figure 4D). These results are consistent with CRX binding to motifs within a broader context than just sequence. Previously, we had shown the effects of DNA methylation on the structure of DNA and the CRX binding motif ^26^. In addition, changing the base pair sequence alters the local base stacking energies, which affect the structure and dynamics of the DNA molecule ^32–34^. Thus, within the broader context of the DNA molecule in the assay, the second site may not be the preferred binding site for CRX-DBD or the base substitution does not disrupt the contacts between the CRX-DBD and DNA under the assay conditions. Thus CRX binding motifs may be a marker of potential CRX binding locations but there are additional effects that dictate CRX specificity.

Outside of the CRX-DBD, the activation domain, which is ∼60% of the total protein length, is predicted by many approaches to be unstructured (Figure 1B). When we plotted the known types of pathogenic mutations from the ClinVar repository at each amino acid position, we observed that the single amino acid substitutions in the activation domain linked to disease are less prevalent than frameshift changes in this region or introduction of a premature stop codon (Figure 5). In contrast, the single amino acid substitutions are more likely to be pathogenic in the DBD. Other studies of disease-linked amino acid substitutions have noted that frameshift mutations in the activation domain lead to severe disease, but they have not identified significant numbers of point mutants in this region ^35–39^. This information supports that the activation domain does not have a specific sequence associated with the structure and requires large changes in sequence to induce effects on protein function. In support of this idea, Chen and coworkers showed that CRX constructs with single amino acid substitutions in the activation domain showed near-normal transactivation, while frameshifts and early stops in the activation domain strongly affected activity ^31^. These clinical findings support a region with a dynamic structure. A disordered activation domain seems to be at odds with the number of proteins that are known or predicted to interact with CRX in this region ^40^. However, flexible binding sites within proteins are recognized with increasing frequency as important to cellular processes, including transcription ^41–43^. Given the significance of the activation domain region to gene regulation by CRX, the form and function of the CRX activation domains warrant further study.

**Figure 5:**
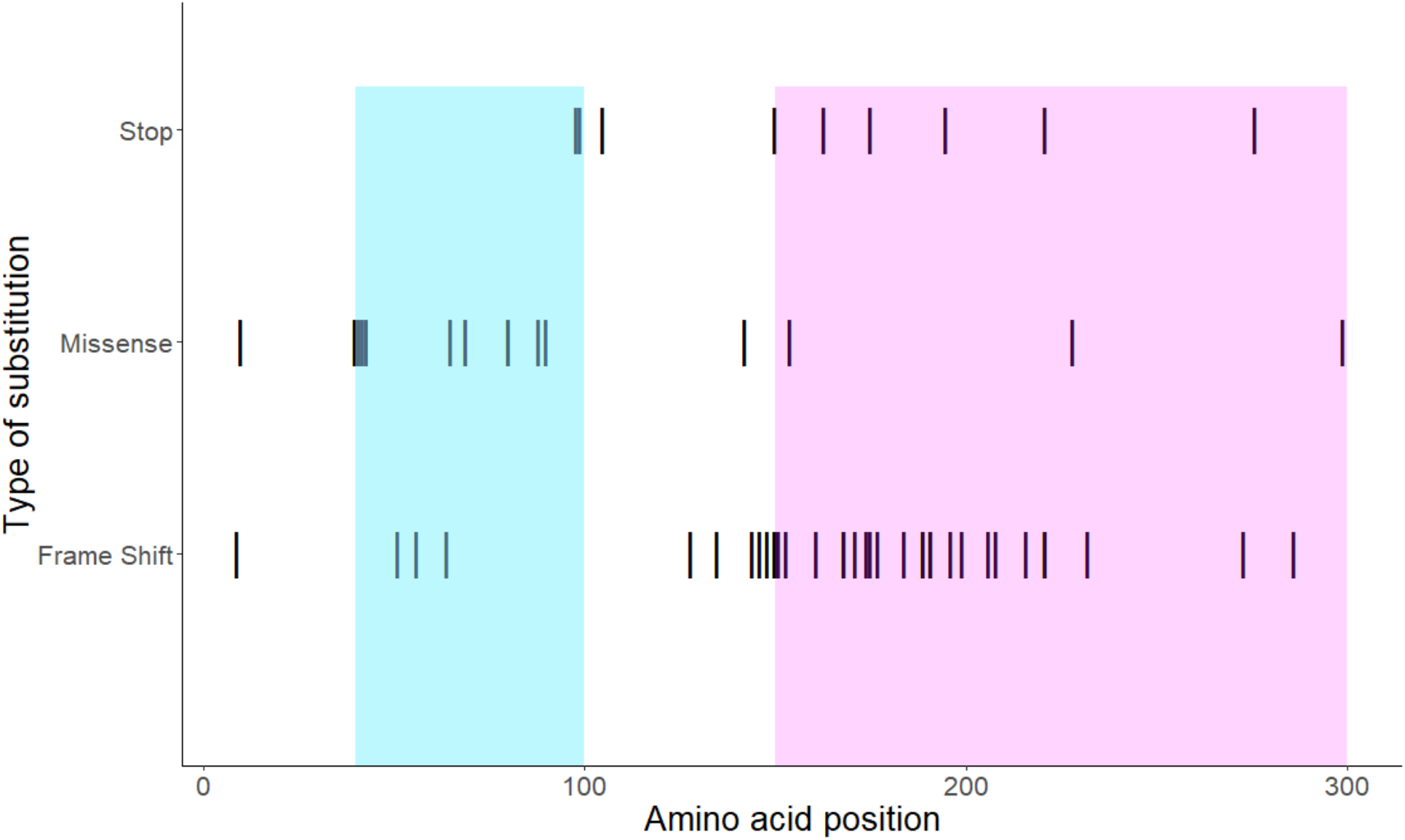
Distribution of clinically relevant mutations in the human CRX protein. Stop, missense, and frameshift mutations in human CRX protein were obtained from the ClinVar database. The distribution of mutations was plotted by amino acid position in the human CRX protein with cyan indicating the DNA binding domain and salmon box indicating the activation domain. The DBD contains 13 pathogenic missense mutations and 3 pathogenic frameshift mutations. The activation domain contains 3 pathogenic missense mutations and 25 pathogenic frameshift mutations.

## Acknowledgments

This work was supported in part by award NIH NEI R15EY028725 (to R.E.), the 4-VA organization, and equipment purchased by an award from the Thomas and Kate F. Jeffress Memorial Trust (to C.E.B.). SAXS data were collected at SIBYLS which is supported by the DOE-BER IDAT DE-AC02-05CH11231 and NIGMS ALS-ENABLE (P30 GM124169 and S10OD018483).

## Data Availability Statement

The SAXS data that support the findings of this study are openly available in SASBDB at https://www.sasbdb.org/ under reference number SASDMA5

## Conflict of Interest Statement

The authors have no conflicts of interest to declare.

**Table 1:** SAXS data collection and analysis parameters.

|  |  |
| --- | --- |
| **Instrument** | SIBYLS beamline 12.3.1 |
| **Wavelength (Å)** | 1.03 |
| **$q$ range (Å<sup>-1</sup>)** | 0.027-0.46 |
| **Injected Concentration (mg mL<sup>-1</sup>)** | 7 |
| **Temperature (K)** | 283 |
| **Column** | Shodex KW-802.5 |
| **$I(0)$ (A.U.) [from $p(r)$]** | 0.93±0.01 |
| **$R_g$ (Å) [from $p(r)$]** | 12.4±0.1 |
| **$I(0)$ (A.U.) [from Guinier]** | 0.91±0.01 |
| **$R_g$ (Å) [from Guinier]** | 11.8±0.2 |
| **$D$ max (Å)** | 40.0 |
| **Porod volume estimate (Å<sup>3</sup>)** | 3670 |
| **Molecular mass Bayes (Da)** | 5700 [99.6% C.I. 5.0 to 6.0] |
| **Calculated monomeric $M_r$ from the sequence (Da)** | 7369 |
| **Primary data reduction** | Scatter 4.0d and RAW 2.1.1 |
| **Data processing** | RAW 2.1.1 |
| **Dummy Atom modeling** | DAMMIF |
| **Electron Density calculation** | DENSS |
| **Three-dimensional graphic representations** | Chimera |

